# GLT8D1 mutations cause amyotrophic lateral sclerosis via disruption of neurotrophin signalling within membrane lipid rafts

**DOI:** 10.1101/2022.06.28.497990

**Authors:** Tobias Moll, Emily Graves, Agnieszka Urbanek, Nikita Soni, Ramya Ranganathan, Adrian Higginbottom, Shanshan Wang, Brian P Head, Johnathan Cooper-Knock, Pamela J Shaw

**Affiliations:** Sheffield Institute for Translational Neuroscience, University of Sheffield, Sheffield, UK; University of California San Diego, California, USA

**Keywords:** GLT8D1, CAV1, amyotrophic lateral sclerosis, Golgi, membrane lipid rafts, gangliosides, neurotrophin signalling

## Abstract

Mutations within *GLT8D1* contribute to familial amyotrophic lateral sclerosis. Pathogenic mutations impair GLT8D1 glycosyltransferase enzymatic function via a dominant negative mechanism, yet the downstream mechanism leading to neurotoxicity is unclear. Here we show that a p.R92C mutation causes fragmentation of the Golgi network and reduces ganglioside expression within membrane lipid rafts (MLRs), leading to impaired neurotrophin signalling. Expression of p.R92C-GLT8D1 in HEK293 cells and mouse primary neurons reduces expression of GM1 gangliosides within the cell plasma membrane leading to disruption of MLRs. Furthermore, p.R92C-GLT8D1 reduces TrkB-mediated pro-survival signalling in MLRs isolated from primary neurons. Interestingly, up-regulation of wild-type GLT8D1 enhances MLRs and promotes pro-survival signalling through TrkB. This closely mirrors findings for another ALS gene, *CAV1*, suggesting convergence on a common pathogenic pathway. Other ALS genes have been associated with Golgi dysfunction and may disrupt the same pathway, suggesting a potential new therapeutic approach via upregulation of GLT8D1.

**Graphical Abstract:** 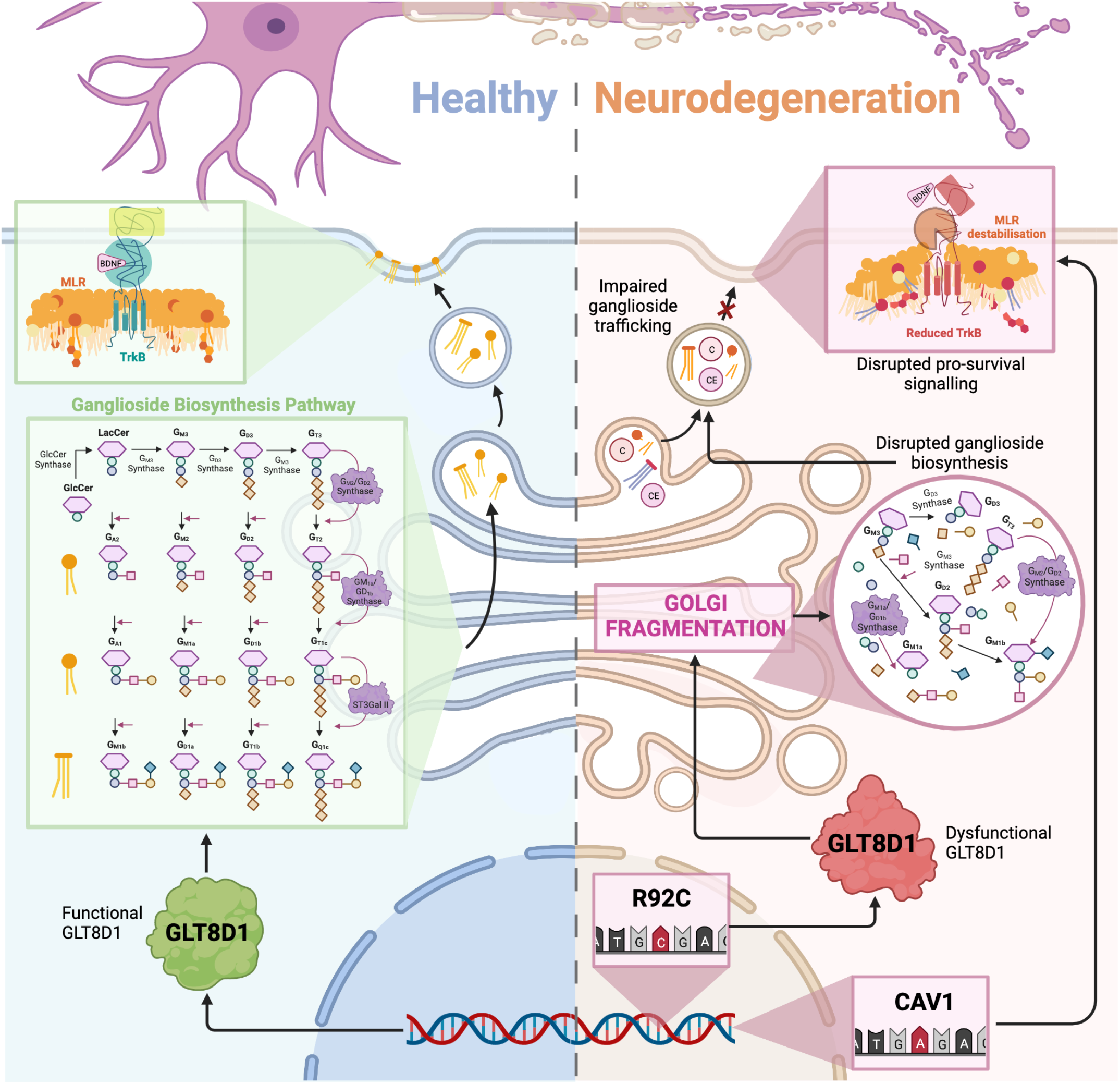

## Introduction

Amyotrophic lateral sclerosis (ALS) is a relatively common neurodegenerative disorder characterised by the loss of motor neurons leading to muscle atrophy, paralysis, and death usually by respiratory failure within 2-3 years of symptom onset (Hardiman et al., 2017). Ten percent of ALS is monogenic, most commonly associated with mutations in either *SOD1* (Rosen et al., 1993) or *C9ORF72* (DeJesus-Hernandez et al., 2011; Renton et al., 2011). Ninety percent of ALS is thought to result from a complex interaction of a polygenic risk genotype and environmental modifiers. Convergence of ALS risk genes within biological pathways has defined current understanding of pathogenesis. To date, these include failure to clear toxic protein aggregates (Aladesuyi Arogundade et al., 2019; Haeusler et al., 2014), aberrant RNA processing leading to neurotoxic dipeptides (Donnelly et al., 2013; Gendron et al., 2013; Mizielinska et al., 2013), oxidative stress (Obrador et al., 2020), mitochondrial dysfunction (Nakaya & Maragkakis, 2018), and axonal transport defects (Burk & Pasterkamp, 2019).

We previously described fully penetrant ALS-associated mutations within the gene encoding the glycosyltransferase GLT8D1 (Cooper-Knock et al., 2019). These findings have since been corroborated in a different population (Tsai et al., 2021). Discovered mutations are proximal to the substrate binding domain and impair GLT8D1 enzyme activity. Glycosyltransferases function prominently in the biosynthesis of gangliosides, which are sialic acid-containing glycosphingolipids expressed abundantly within the central nervous system (CNS) (Moll et al., 2020). Gangliosides within the CNS are synthesized in the endoplasmic reticulum (ER) from a lactosylceramide precursor and are remodelled during transit from the *cis*-to the *trans*-Golgi network by a series of glycosyltransferase enzymes that incorporate galactose and N-acetylgalactosamine groups (Moll et al., 2020). Consistent with a role in this process, GLT8D1 shows prominent CNS expression, carries a Golgi localisation signal, and accepts galactose as a substrate (Cooper-Knock et al., 2019).

Mature gangliosides are transported to the cell plasma membrane by vesiculation where they provide structure and stability to membrane lipid rafts (MLRs), specialised plasma membrane microdomains that regulate a wide range of cellular physiology including signal transduction, spatial organisation of the plasma membrane, and endocytosis (Grassi et al., 2020; Koyama-Honda et al., 2020; van IJzendoorn et al., 2020). In neurons, MLRs serve as platforms for pro-survival and pro-growth signalling cascades initiated by neurotrophic factors (e.g., BDNF/TrkB activation) (Moll et al., 2021; Pike, 2005; Sawada et al., 2019). Disrupted MLRs are associated with impaired neurotrophin signalling and consequent neurodegeneration (Sawada et al., 2019). Dysregulated neurotrophin signalling is a well described mechanism in the context of ALS (Pradhan et al., 2019) and in the field of neurodegeneration more broadly (Mitre et al., 2017).

Another important constituent of MLRs is the cholesterol binding protein, caveolin-1 (CAV1), which forms a hetero-oligomeric complex with caveolin-2 (CAV2) to facilitate the organisation and targeting of neurotrophic receptor signalling pathways (de Almeida, 2017; Head et al., 2011; Head et al., 2008; Sawada et al., 2019). CAV1 upregulation enhances MLRs and promotes neurotrophin signalling, leading to enhanced neuronal survival (Head et al., 2011; Mandyam et al., 2017). In contrast, loss of CAV1 accelerates neurodegeneration (Head et al., 2011; Head et al., 2010). Neuron-specific upregulation of CAV1 is neuroprotective and improves motor and cognitive function in pre-clinical models of ALS and Alzheimer’s disease respectively (Egawa et al., 2017; Sawada et al., 2019; Wang et al., 2021). We have previously described ALS-associated genetic variation within enhancer elements linked to the expression of CAV1 and CAV2. These enhancer mutations reduce CAV1/CAV2 expression and disrupt MLRs in patient-derived cells (Cooper-Knock et al., 2020).

We hypothesised that ALS-associated mutations leading to impaired function of *GLT8D1* and *CAV1* may have a common impact on the neurotrophin signalling cascade via MLRs. To test this, we investigated the effect of the most common and clinically aggressive mutation (p.R92C) on membrane ganglioside expression in HEK293 cells and mouse primary neurons. In both models we observed mutation-specific reductions in membrane ganglioside expression leading to disruption of MLRs. We subsequently isolated MLRs from primary neurons overexpressing wild-type (WT) or mutant (p.R92C) forms of GLT8D1 by sucrose density fractionation. In MLRs, the neurotrophic receptor, TrkB, was reduced in mutant cells suggesting impaired pro-survival signalling. To understand why GLT8D1 mutations might cause destabilisation of MLRs, we investigated Golgi fragmentation as a potential explanation for impaired ganglioside trafficking. We observed fragmentation of the *cis*-Golgi cisternae in HEK293 cells following stable expression of the p.R92C-GLT8D1 mutant protein compared to WT and isogenic controls. This was validated in primary neurons following lentiviral transduction of WT- and p.R92C-mutant *GLT8D1*. Based on these lines of evidence we argue that both CAV1 and GLT8D1 converge on a similar mechanism of pathogenesis in ALS: impaired neurotrophin signalling via disruption of MLRs.

## Results

### ALS-associated p.R92C-GLT8D1 mutation disrupts MLRs

MLRs are enriched with gangliosides that promote a stable microenvironment for the modulation of cell signalling (Grassi et al., 2020; Schengrund, 2015). Given that ganglioside expression is directly linked to glycosyltransferase activity (Grassi et al., 2020; Moll et al., 2020; Ngamukote et al., 2007), we hypothesised that the ALS-associated p.R92C mutation within GLT8D1 would disrupt membrane ganglioside expression and, by extension, MLR integrity.

Initially, we induced stable expression of either WT-GLT8D1 or p.R92C-GLT8D1 in HEK293 cells (**Figure S1, Methods**) to investigate ganglioside expression by live-cell imaging. Sialic acid and N-acetylglucosaminyl residues, which are important constituents of gangliosides (Schnaar et al., 2014), were labelled using a wheat germ agglutinin (WGA) molecular probe (Chazotte, 2011). Whole membrane fluorescence intensity of WGA was markedly reduced in p.R92C mutant cells compared to WT cells (20% reduction, *p* = 0.0145, paired *t*-test, **Figure 1A-B**), suggesting disruption of membrane ganglioside expression. To further characterise the effect of the p.R92C mutation on lipid raft integrity we used the MLR-specific probe, CT-B, which binds with high affinity to ganglioside GM1 (Aman et al., 2001; Day & Kenworthy, 2015; Head et al., 2011; Sawada et al., 2019). We observed a reduction in membrane fluorescence intensity of CT-B in HEK293 cells following stable expression of p.R92C-GLT8D1 compared to stable expression of WT-GLT8D1 (10% reduction, *p* = 0.0277, paired *t*-test, **Figure 1C-D**).

**Figure 1:**
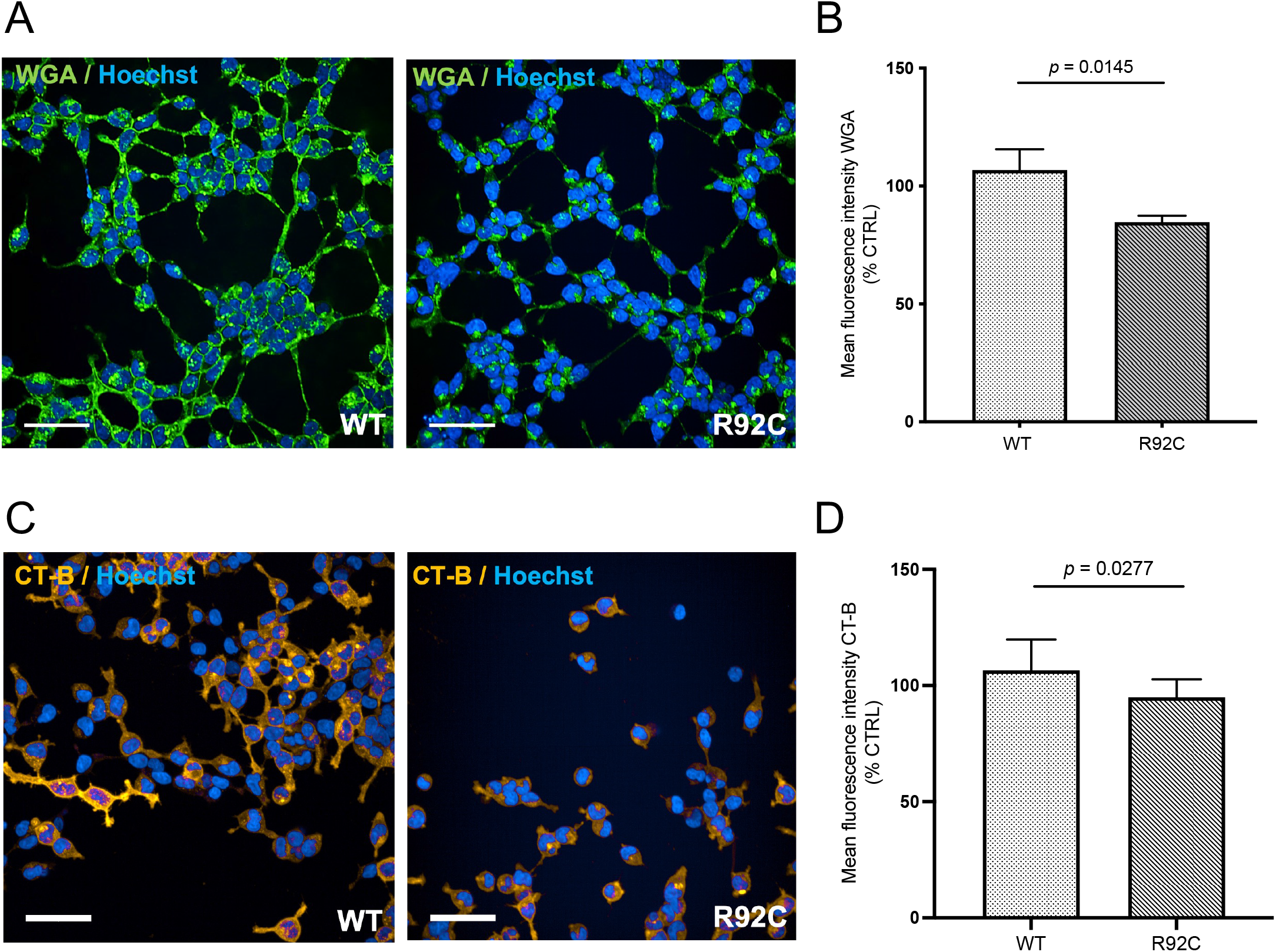
ALS-linked p.R92C-GLT8D1 mutation disrupts MLRs in HEK293 cells. (**A**) Representative staining of sialic acid residues in HEK293 cells stably overexpressing WT-*GLT8D1* or p.R92C-*GLT8D1*. Sialic acids were labelled using a WGA molecular probe (green). (**B**) Fluorescence intensity of membrane sialic acid residues is reduced by ∼20% in mutant compared to WT. (**C**) Representative staining of MLRs in WT and mutant cells detected using CT-B molecular probe (orange). (**D**) Fluorescence intensity of membrane CT-B is reduced in mutant compared to WT (∼10% reduction). Nuclear counterstain (Hoechst 33342) is shown in blue. Scale bars = 50μM. Each data point is expressed as a percentage of control (sham-transfected) (paired *t*-test, *p* = 0.0277). Error bars represent mean ±SD. Images are enhanced for presentation only. Quantifications were performed on raw image data.

We corroborated our findings in an ALS-relevant cell model through transduction of mouse primary neurons with WT-GLT8D1 or p.R92C-GLT8D1 lentiviral constructs (**Methods**). Transduction efficiency was determined according to GFP expression (**Figure S2**). Membrane CT-B fluorescence intensity was comparably reduced in primary neurons transduced with p.R92C-GLT8D1 compared to WT-GLT8D1, suggesting not only a role for GLT8D1 in neuronal ganglioside biosynthesis but also a mutation-specific effect on MLR integrity (7% reduction, *p* = 0.04, unpaired *t*-test, **Figure 2A-B**). Our findings place dysregulated ganglioside expression and MLR integrity downstream of GLT8D1 enzyme dysfunction.

**Figure 2:**
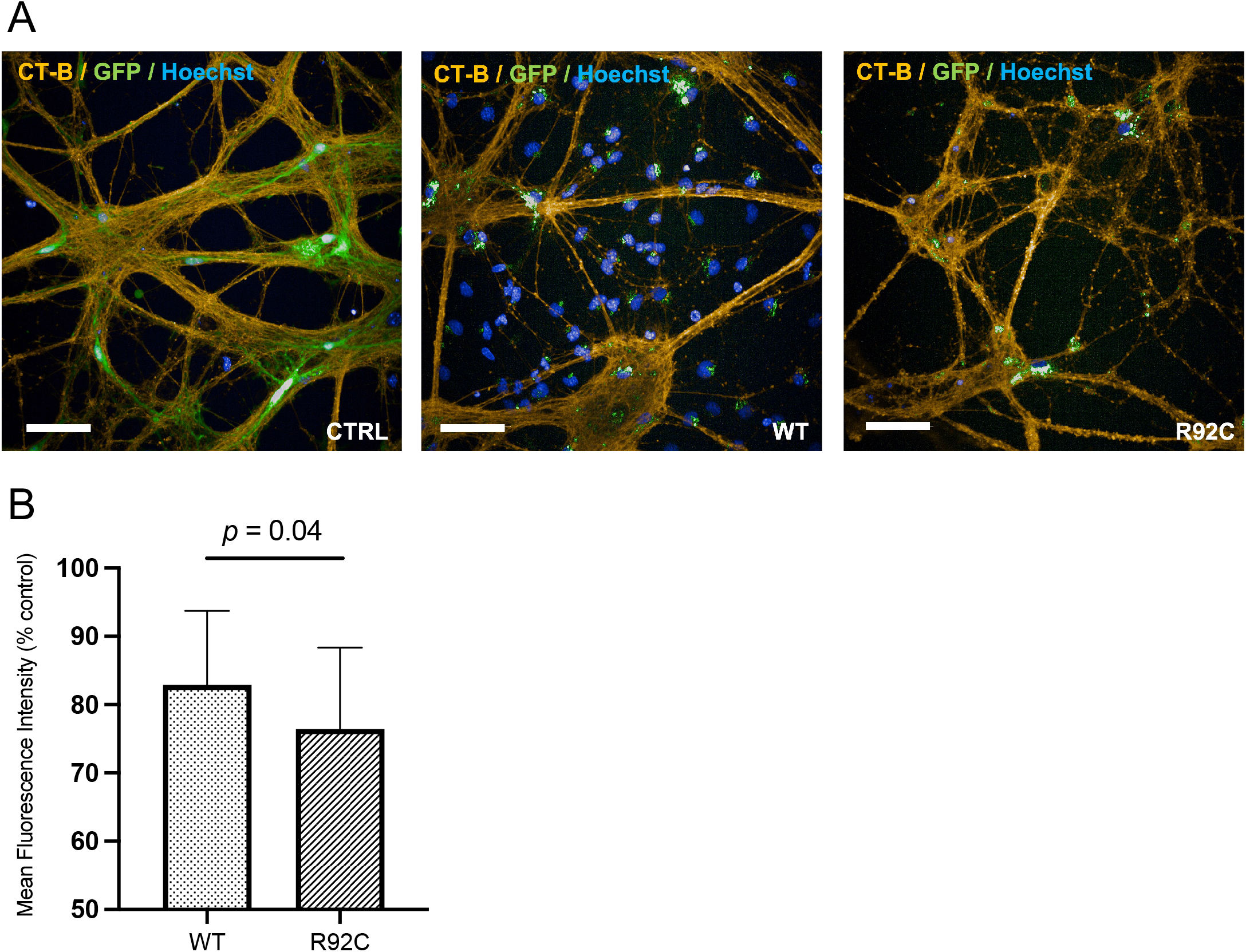
ALS-associated p.R92C-GLT8D1 mutation disrupts MLRs in mouse primary neurons. (**A**) Representative images of MLRs in mouse wild-type primary neurons transduced with either LV_GLT8D1[WT]-eGFP, LV_GLT8D1[p.R92C]-eGFP, or LV-eGFP. CT-B (orange) was used as a marker of MLRs; nuclear counterstain (Hoechst 33342) is shown in blue. (**B**) Mean fluorescence intensity of CT-B is reduced in primary neurons transduced with LV_GLT8D1[p.R92C]-eGFP compared to LV_GLT8D1[WT]-eGFP (*p* = 0.04, unpaired *t*-test). Data are expressed as percentage of cells transduced with LV-eGFP. Scale bars = 50μM. Error bars represent mean ±SD. Images are enhanced for presentation only. Quantifications were performed on raw image data.

### ALS-associated p.R92C-GLT8D1 mutation causes fragmentation of the Golgi network

Next, we investigated Golgi fragmentation within models expressing p.R92C-GLT8D1. Golgi fragmentation is a well-described feature of neurodegenerative diseases including ALS and has been shown to occur upstream of motor neuron loss (Haase & Rabouille, 2015; Maruyama et al., 2010; Mourelatos et al., 1996; van Dis et al., 2014). The observed reductions in membrane ganglioside expression may be consistent with dysregulation of Golgi trafficking. GLT8D1 has a Golgi localisation signal that is unaffected by the p.R92C mutation (Cooper-Knock et al., 2019). Moreover, glycosyltransferases have previously been shown to influence Golgi architecture (Nilsson et al., 2009). We hypothesised that the ALS-associated p.R92C mutation within GLT8D1 would cause fragmentation of the Golgi network.

In HEK293 cells stably expressing either WT-GLT8D1 or p.R92C-GLT8D1, we observed mutation-specific fragmentation of the *cis*-Golgi cisternae in p.R92C-mutant cells compared to WT cells and isogenic controls (*p* = 0.0383, one-way ANOVA, **Figure 3A-B**). We observed a trend towards increased fragmentation of the *trans*-Golgi (*p* < 0.05, one-way ANOVA, **Figure 3C-D**) cisternae in p.R92C-mutant cells compared to WT cells and isogenic controls, although this was not statistically significant. We confirmed fragmentation of the *cis*-Golgi cisternae in mouse primary neurons transduced with p.R92C-GLT8D1 lentivirus compared to WT-GLT8D1 (*p* = 0.0478, unpaired *t*-test, **Figure 3E-F**). Based on these data, it is possible that Golgi fragmentation precedes MLR instability in GLT8D1-ALS.

**Figure 3.**
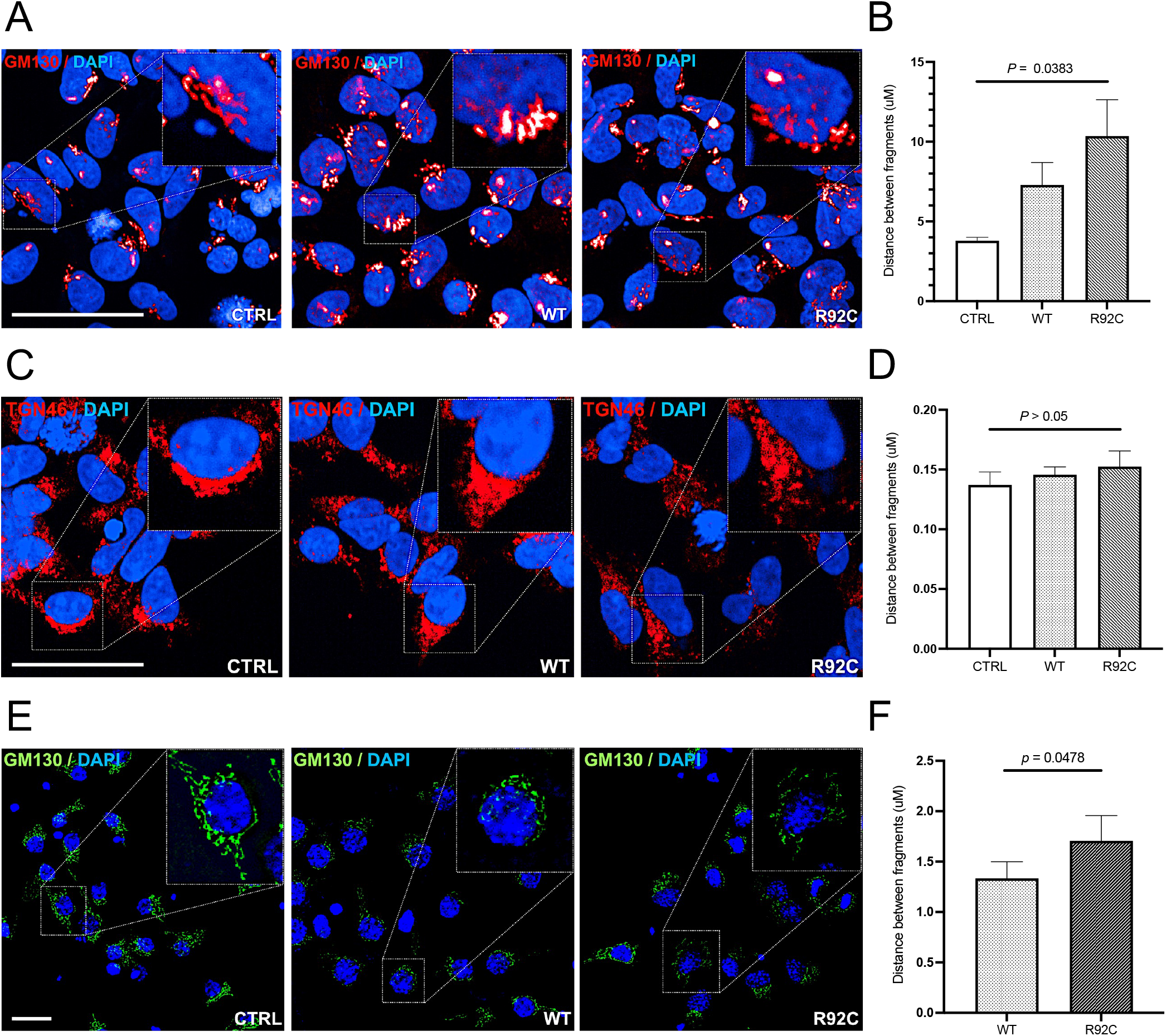
Overexpression of p.R92C-GLT8D1 causes fragmentation of the Golgi network in HEK293 cells. Representative staining of the (**A**) *cis*-Golgi and (**C**) *trans*-Golgi cisternae in CTRL (GFP-only), WT-GLT8D1, and p.R92C-GLT8D1 HEK293 cells and (**E**) primary neurons. The image segments outlined with a white box are magnified to delineate Golgi morphology; Golgi markers GM130 (*cis*-Golgi) and TGN46 (*trans*-Golgi) are shown in red. (**B, D, F**) The distance between the nearest neighbouring fragment (μM) was measured for each condition [(HEK293) GM130: one-way ANOVA with Welch’s correction, *p* = 0.0383; TGN46: one-way ANOVA with Welch’s correction, *p* > 0.05]; [(primary neurons) GM130: *p* = 0.0478, unpaired *t*-test]. Nuclear counterstain (Hoechst 33342) is shown in blue. Scale bars = 50μM. Error bars represent mean ±SD. Images are enhanced for presentation only. Quantifications were performed on raw image data.

### ALS-associated p.R92C-GLT8D1 mutation disrupts neurotrophin signalling within MLRs

In neurons, pro-survival signalling cascades occur predominantly through neurotrophin receptors associated with MLRs (Chiricozzi et al., 2020; Egawa et al., 2017; Moll et al., 2021; Zonta & Minichiello, 2013). These cascades are mediated through tropomyosin receptor kinases (Trk) and p75 neurotrophin receptors (p75^NTR^) (Moll et al., 2021). The localisation of neurotrophin receptors to MLRs is regulated by gangliosides, and multiple lines of evidence demonstrate that the ganglioside GM1 is essential for the activation of Trk receptors (Bachis et al., 2002; Duchemin et al., 2002; Farooqui et al., 1997; Mutoh et al., 1995; Pitto et al., 1998; Rabin et al., 2002). In the present study, we observed a reduction in membrane GM1 expression in neuronal and non-neuronal cells expressing an ALS-associated p.R92C mutation within *GLT8D1*. We therefore hypothesised that loss of membrane GM1 would negatively impact Trk expression within MLRs, disrupting neurotrophin signalling leading to subsequent neurodegeneration.

We isolated MLRs from primary neurons overexpressing WT-*GLT8D1* or p.R92C-*GLT8D1* by sucrose density fractionation (Head et al., 2011) (**Methods**). Successful isolation of lipid rafts within buoyant fractions 4 and 5 was confirmed by immunoblotting for CT-B and CAV2 (**Figure S3**). Immunoblotting within these fractions revealed a 10-15% reduction in TrkB expression in lipid rafts isolated from mouse primary neurons expressing the p.R92C-*GLT8D1* mutant protein compared to WT-*GLT8D1* and isogenic controls (**Figure 4A-B, Methods**). Interestingly, MLR integrity, as measured by CT-B expression, was enhanced by >200% in cells overexpressing WT-*GLT8D1* (**Figure 4A, C**).

**Figure 4:**
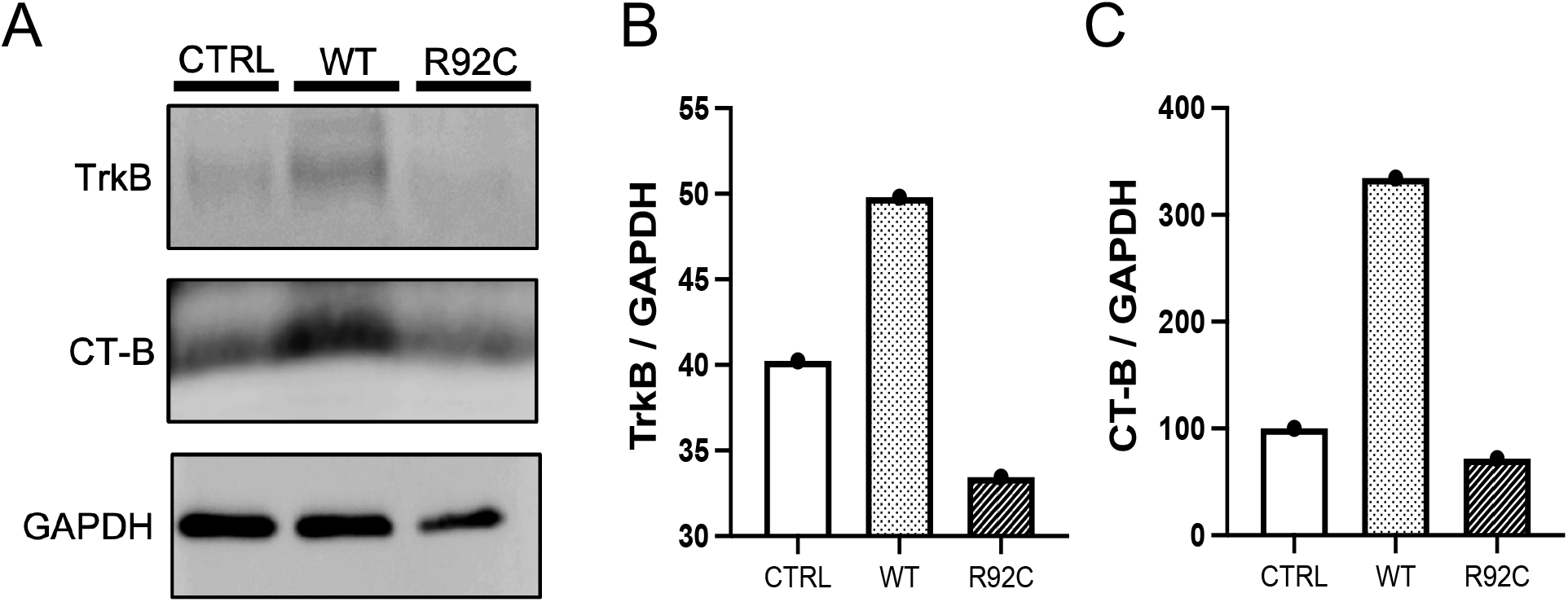
GLT8D1 stabilizes MLRs and enhances the expression of neurotrophin signaling components in primary neurons. (**A**) Immunoblots showing increased intensity of TrkB and CT-B (lipid raft marker) in primary neurons transduced with either LV_GLT8D1[WT]-eGFP, LV_GLT8D1[R92C]-eGFP, or LV-eGFP. TrkB expression is enhanced in cells transduced with LV_GLT8D1[WT]-eGFP but reduced in cells transduced with LV_GLT8D1[R92C]-eGFP. (**B**) Densitometry data are expressed as a percentage of GAPDH (n=1).

## Discussion

ALS genes are noted to cluster within certain biological pathways including RNA processing, axonal transport, and protein homeostasis (Cooper-Knock et al., 2021; Mejzini et al., 2019). We have demonstrated that ALS-associated mutations within GLT8D1 disrupt MLRs and neurotrophin signalling, which closely mirrors previous findings for CAV1-ALS (Head et al., 2011; Mandyam et al., 2017), thereby adding a new biological pathway with convergent genetic risk. The link between ALS genetics and the disruption of MLRs is not new. For example, alterations in the MLR proteome have previously been described in the SOD1^G93A^ mouse model of ALS (Zhai et al., 2009), and loss of the ALS receptor protein, sigma-1, destabilises MLRs (Al-Saif et al., 2011; Mavlyutov et al., 2013; Vollrath et al., 2014). However, association of this pathway with genetic changes, which are present at conception, necessarily places MLR dysfunction upstream in the pathogenesis of a late age-of-onset disease such as ALS.

Glycosyltransferase dysfunction is linked to the pathogenesis of neurodegenerative disorders including ALS (Moll et al., 2020). Glycosyltransferases exhibit diverse functions and for many of these enzymes the biological pathway associated with their activity is unclear. However, glycosyltransferase dysfunction associated with neurodegenerative disease appears to converge on ganglioside metabolism (Moll et al., 2020). Indeed, gangliosides have been shown to modulate ALS pathogenesis (Dodge et al., 2015). In the present study we describe a mutation-specific reduction in membrane ganglioside expression. In the context of ALS, changes in ganglioside expression patterns have been reported in both directions. For example, SOD1^G93A^ mice show reduced ganglioside levels in motor axons in both early symptomatic and end stages of the disease (Dodge et al., 2015; Henriques et al., 2017); in contrast gangliosides were shown to be increased in ALS patient post-mortem tissue (Rapport et al., 1985). Our findings demonstrate that dysregulated ganglioside metabolism upstream of the pathogenesis of *GLT8D1*-ALS, and provides justifiable cause to investigate gangliosides, and GM1 in particular, as potential therapeutic targets in patients with ALS-linked *GLT8D1* mutations. Indeed, gangliosides have already been explored for decades as potential therapeutic targets in a variety of neurodegenerative disorders including ALS for several decades, although earlier clinical studies lacked experimental power (Bradley, 1984; Hallett et al., 1984; Harrington et al., 1984; Henriques et al., 2017; Knight et al., 2015; Schneider et al., 2015).

Ganglioside biosynthesis is compartmentalised within the Golgi apparatus and is organised in distinct units formed by associations with glycosyltransferases (Giraudo & Maccioni, 2003). We have demonstrated that ALS-associated *GLT8D1* mutations are associated with both reduced ganglioside synthesis and fragmentation of the Golgi network, which we link to MLR instability and deficient neurotrophin signalling. Golgi fragmentation has been identified in up to 70% of familial ALS patient motor neurons (Fujita et al., 2000; Ito et al., 2011) and 10-50% of sporadic ALS patient motor neurons (Gonatas et al., 2006; van Dis et al., 2014) bearing *SOD1, FUS* or *OPTN* mutations. *CAV1* has also been shown to influence Golgi architecture (Tiwari et al., 2016). Precedent even exists for a link between the cytoplasmic mislocalisation of TDP-43, the pathological hallmark of most ALS cases, and Golgi fragmentation (Fujita et al., 2008). Golgi fragmentation has been shown to occur prior to neuromuscular denervation, suggesting that it occurs upstream in the pathophysiology of ALS (van Dis et al., 2014). It is therefore possible that ganglioside synthesis, MLR function, and neurotrophin signalling are impaired more broadly in ALS, outside of mutations associated with *GLT8D1* and *CAV1*.

Gangliosides are important constituents of MLRs, which have a key role in the organisation of cell signalling and neuronal survival. Impaired neurotrophin signalling pathways, particularly the TrkB signalling pathway, are a well described feature in ALS pathophysiology (Bronfman et al., 2007; Pradhan et al., 2019; Sleigh et al., 2019), which reflects our finding here that ALS-associated *GLT8D1* mutations impair TrkB signalling in mouse primary neurons. Brain-derived neurotrophic factor (BDNF) binds to TrkB receptors to promote pro-growth and pro-survival signalling. However, the neurotrophic activity of TrkB is directly modulated through its interaction with endogenous GM1 within the membrane (Choucry et al., 2019; Duchemin et al., 2002; Ledeen & Wu, 2015; Pitto et al., 1998). In the present study overexpression of WT-GLT8D1 led to an up-regulation of membrane GM1, which was accompanied by increased expression of TrkB. It is important to note that neurotoxicity resulting from TrkB overactivation has been reported in the context of ALS (Fryer et al., 2000; Hu & Kalb, 2003; Kust et al., 2002). In addition, there have been several reports describing the expression of anti-ganglioside antibodies in people with motor neuropathies (Annunziata et al., 1995; Kornberg, 2000; Mizutani et al., 2003; Pestronk & Choksi, 1997; Stevens et al., 1993). Studies such as these highlight the importance of maintaining a finely balanced and tightly regulated microenvironment homeostasis to ensure proper motor neuron function.

Taken together, our results suggest that the homeostasis of gangliosides is dysregulated in *GLT8D1*-ALS. Loss of ganglioside homeostasis appears to destabilise MLRs causing disrupted neurotrophin signalling. Impaired neurotrophin signalling through MLRs is a well-described feature of ALS pathophysiology (Head et al., 2011; Moll et al., 2021), and targeted re-stabilisation of MLRs via up-regulation of CAV1 has clear therapeutic potential (Egawa et al., 2017; Sawada et al., 2019; Wang et al., 2021). We have previously identified ALS-associated genetic variation within enhancer elements linked to the expression of CAV1. In the context of ALS, both GLT8D1 and CAV1 are associated with a loss of MLR stability. The precise regulation of these highly dynamic microdomains is essential for maintaining neuronal health and preventing neurodegeneration (Schengrund, 2010). We propose that both CAV1 and GLT8D1 converge on a similar pathway: impaired neurotrophin signalling due to disruption of MLRs. Selective targeting of these proteins has the potential for therapeutic benefit in certain genetic subtypes of ALS.

## Figure legends

**Supplementary Figure S1:**
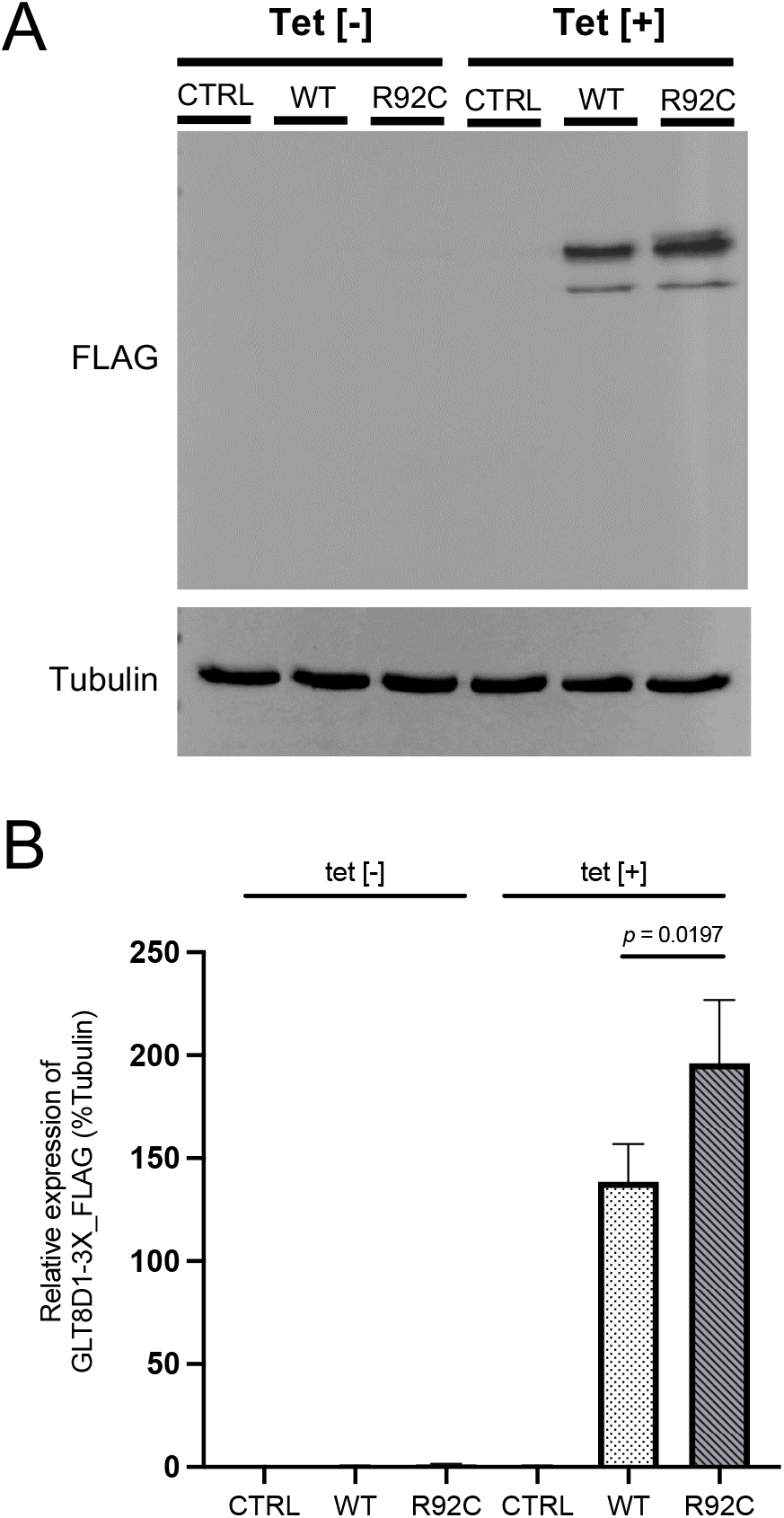
(**A**) Western blot showing tetracycline-induced expression of WT-GLT8D1 and p.R92C-GLT8D1 in isogenic HEK293 cells. Relative GLT8D1 expression was detected using an anti-FLAG antibody; α-Tubulin was used as a loading control. (**B**) Densitometric analysis shows enhanced levels of p.R92C-GLT8D1 compared to WT-GLT8D1 (paired *t*-test, p = 0.0145).

**Supplementary Figure S2:**
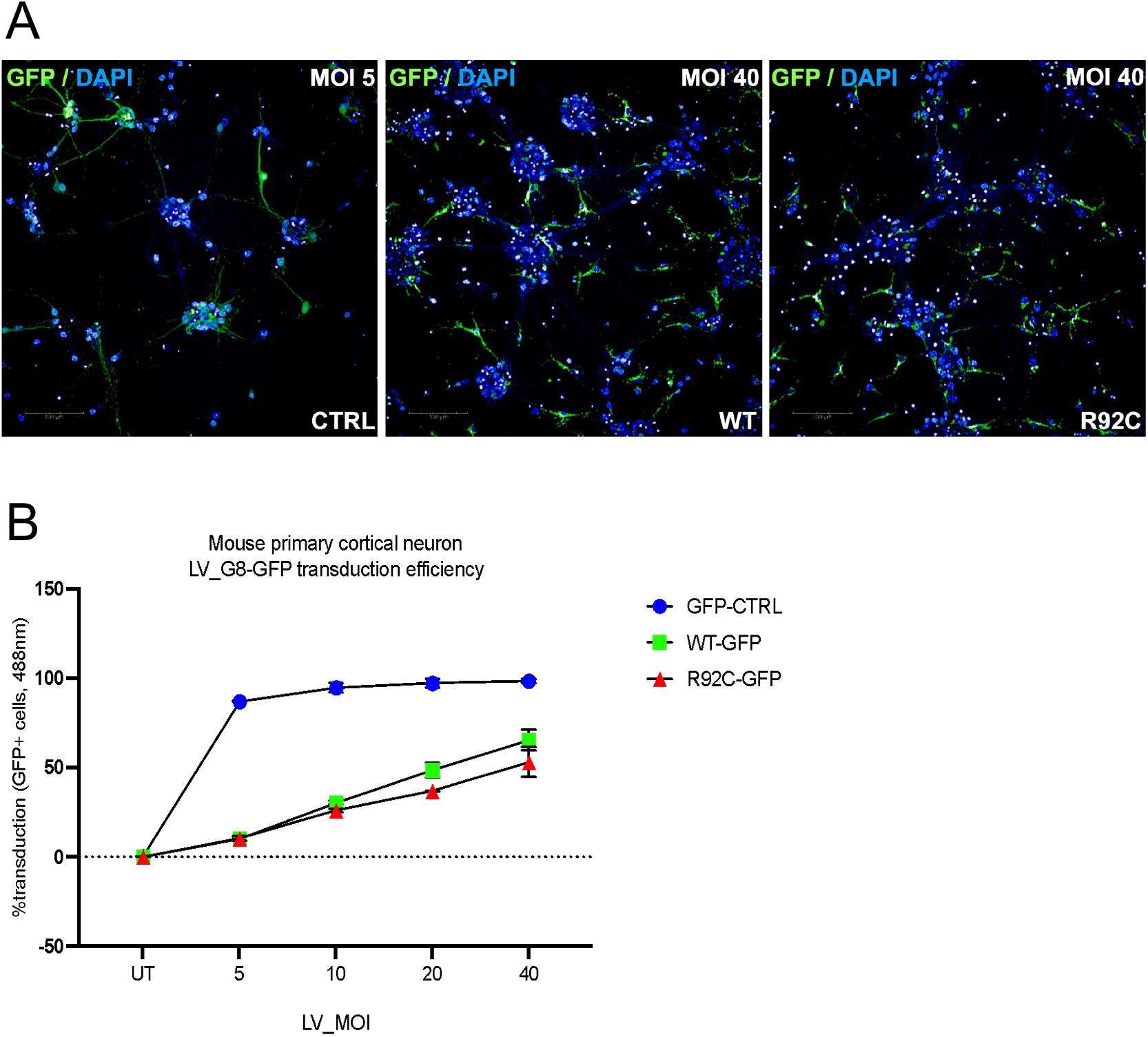
(**A**) Representative images of GFP expression in mouse primary neurons transduced with GFP-only, WT-GLT8D1 or p.R92C-GLT8D1 lentivirus. (**B**) MOI was determined by the number of GFP-positive cells. For experimental work, a MOI of 40 was used for WT-GLT8D1 and p.R92C-GLT8D1, and a MOI of 5 was used for GFP-only control. MOI 40 produced ∼50% transduction efficiency across both GLT8D1 cell populations.

**Supplementary Figure S3:**
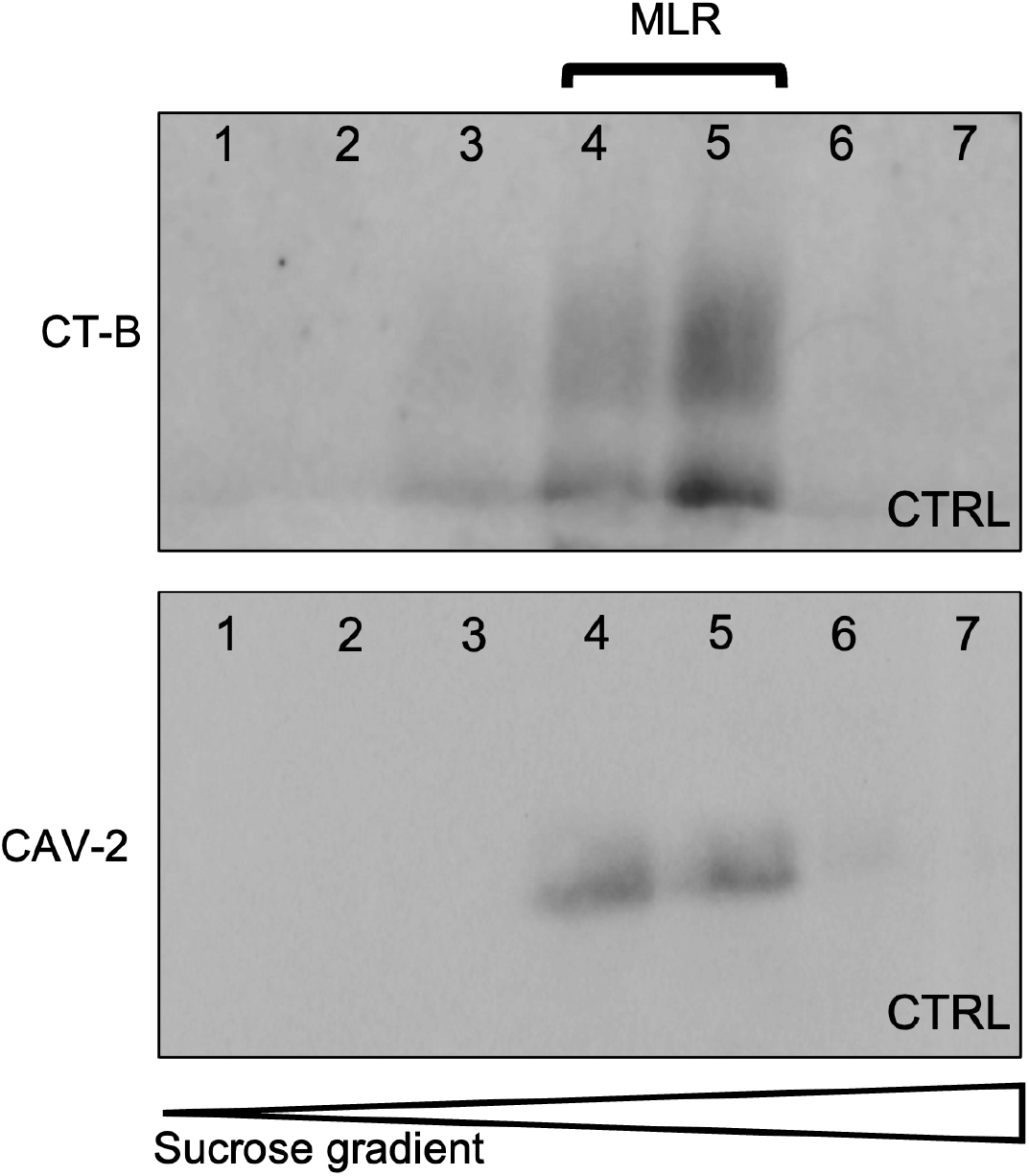
Immunoblots showing isolation of lipid rafts in buoyant fractions 4 and 5 as measured by CT-B and CAV2 expression.

## Materials and Methods

### Engineering of HEK293_GLT8D1-WT/R92C stable cell lines

Stable HEK293 cells exhibiting tetracycline-inducible expression of wild type (WT) and mutant (R92C) GLT8D1 were generated using the Flp-In™ T-REx™ system (ThermoFisher Scientific). The pGKFLPobpA vector (Addgene) encoding a recombinase was co-transfected with either pcDNA5/FRT/TO, pcDNA5/FRT/TO_3xFLAG-GLT8D1, or pcDNA5/FRT/TO_3xFLAG-GLT8D1-R92C plasmids into the HEK293 Flp-In™ T-REx™ CVCL_U427 line in a ratio of 4:6 using polyethylenimine (PEI). Resulting clones were screened for Zeocin™ sensitivity, and Blasticidin S/Hygromycin B resistance, to ensure isogenic.

### HEK293_GLT8D1-WT/R92C stable cell line culture

HEK293 Sham and HEK293_GLT8D1-WT/R92C stable cell lines were cultured in 10mL DMEM (Lonza) supplemented with 10% (v/v) tetracycline-free FBS (Lonza), 50 U/mL Penicillin/Streptomycin (Lonza), 100µg/mL Hygromycin B and 15µg/mL Blasticidin S in 10cm plates at 37°C, 5% CO_2_ and split every 3-4 days. To induce expression of the pcDNA5/FRT/TO_3xFLAG-GLT8D1-WT/R92C construct, cells were cultured in Blasticidin-S-free supplemented DMEM for 24 hours, after which, tetracycline was added to the culture media to create a final concentration of 10mg/mL.

### Preparation and maintenance of dissociated primary mouse cortical neurons

Initially, culture plates were coated in an appropriate volume of poly-D-lysine (0.1mg/mL in dH_2_O) overnight at 37°C, 5% CO_2_. C57BL/6 mice were bred at the University of Sheffield Biological Services Unit and females were sacrificed by cervical dislocation. Cerebral cortices were isolated from embryonic day 15 embryos whilst submersed in cold HBSS-/-. Meninges were manually removed, and cortices dissected using surgical forceps, and the tissue was washed 1x in 10mL HBSS-/-, then resuspended in 5mL HBSS. Trypsin was added to a final concentration of 0.05% and incubated for 15 minutes at 37°C to encourage tissue dissociation. 5mL DNAse (10μg/mL DNAse in HBSS+/+) was added for 2 minutes and the supernatant was aspirated. Tissue was resuspended in 1mL triturating solution (1% albumax, 0.5mg/mL trypsin inhibitor, 10μg/mL DNAse in HBSS ^-/-^) and triturated through flame-polished glass Pasteur pipettes with progressively smaller openings to obtain a single cell suspension. Cells were re-suspended in supplemented neurobasal plus medium (ThermoFisher Scientific) [1x B27 Plus supplement (ThermoFisher Scientific), 1x GlutaMax (ThermoFisher Scientific), 50 U/mL Penicillin/Streptomycin] and maintained at 37°C, 5% CO_2_.

### Generation of GLT8D1 lentiviral vectors

The human GLT8D1 open-reading-frame (NM_001278280.2) was cloned into a pLV[Exp]-eGFP lentiviral vector containing a woodchuck hepatitis virus post regulatory element (W) to overexpress *GLT8D1* under a CMV promoter (pLV_SIN-W-CMV-GLT8D1). All plasmids were validated by Sanger sequencing and sequence traces are available upon request. Lentiviral vectors were constructed by VectorBuilder. Vector IDs (GLT8D1-WT: VB210113-1206cvu; R92C: VB210113-1209wnh) can be used to retrieve detailed information about the vectors on vectorbuilder.com.

HEK293T cells were used for lentiviral production, plated at a density of 3 × 10^6^ per 10 cm dish. Cells were transfected using a calcium chloride transfection containing 0.5M calcium chloride (Sigma), 2X HEPES Buffered Saline (Sigma) and four lentiviral component plasmids; pCMV delta 8.2 (13 µg), pRSV-Rev (3 µg), pMD.G (3.75 µg) (Addgene) and pLV_SIN-W-PGK-EXOSC2 (13 µg) (Delgon et al., 2000). Transfection mix added dropwise to each plate and left overnight, with a full media change carried out the following morning. Cells were incubated for a further 48 hours before all media was collected and filtered using a 0.45 µm filter (Sigma). Equally loaded tubes (Beckman) were then spun at 19,000rpm/90 minutes/4°C using an ultracentrifuge and a SW28 hanging rota (Beckman). All supernatant was removed and each viral pellet was resuspended with 300 µl of 1% bovine serum albumin (Tocris Bioscience) in phosphate buffer solution (Sigma). Each tube was incubated on ice for 1 hour and then combined into one homogeneous solution before being aliquoted and stored at -80°C. Viral titres were measured through qPCR against a virus of known biological titre (FACS titration). Genomic DNA was isolated from cells (as described previously) which had been transduced with a serial dilution of virus. Viral genomic integration was measured using WPRE primers: CCCGTACGGCTTTCGTTTTC (fwd) and CAAACACAGAGCACACCACG (rvs).

### Lentiviral transduction

Mouse primary neurons were left to mature for 7 days prior to transduction. Cells were transduced with lentivirus [GFP (MOI 5); WT-GLT8D1 (MOI 40); p.R92C-GLT8D1, (MOI 40)] for 18 hours at 37°C, 5% CO_2_. Lentivirus was aspirated and replaced with supplemented neurobasal plus medium, and the cells were incubated for a further 2 days at 37°C, 5% CO_2._ Transduction efficiency was calculated according to GFP expression and quantified using ImageJ (NIH).

### Sucrose density fractionation

Cells were lysed in high pH lysis buffer (150 mM Na_2_C0_3_, 1 mM EDTA, + protease inhibitor cocktail, pH 11) followed by sonication (Soniprep 150, MSE) at 50% amplitude for 10 seconds (x3) with 30-second incubations on ice in between. Whole cell lysates were loaded into a sucrose gradient containing either 80%, 35% or 5% sucrose in MES-buffered saline (25 mM MES, 150 mM NaCl, 2 mM EDTA, pH 6.5). Lipid rafts were isolated by sucrose density fractionation by ultracentrifugation at 39,000 rpm for 18 hours at 4°C. Buoyant fractions were precipitated in a 20% solution of trichloroacetic acid (Sigma Aldrich) for 12 hours at -20°C. Samples were centrifuged at 17,000 x g for 20 minutes at 4°C, washed in 100% acetone (Fisher Scientific) (x3), and air dried for 30 minutes on ice.

### Cell lysis

Cells were lysed in IP lysis buffer (150mM NaCl, 50mM HEPES, 1mM EDTA, 1mM DTT, 0.5% (v/v) Triton™ X-100, pH 8.0) containing an EDTA-free protease inhibitor cocktail (PIC) (SIGMAFASTTM Sigma-Aldrich) (20µL/mL) for 15 minutes on ice. The lysate was centrifuged at 17,000 xg for 5 minutes at 4°C and the cell debris pellet was discarded. Protein extracts were quantified using Bradford reagent (BioRAD).

### Western blot analysis

Protein extracts were fractionated via SDS-PAGE in Laemmli buffer (277.8mM Tris-HCl; 44.4% (v/v) glycerol; 4.4% SDS; 0.02% bromophenol blue; 355mM 2-mercaptoethanol; pH 6.8), then electrophoretically transferred to nitrocellulose membranes. Membranes were probed using the relevant primary antibody in 5% (w/v) milk/Tris Buffered Saline, with Tween® 20 (TBST) (20mM Tris, 137mM NaCl, 0.2% (v/v) Tween® 20, pH 7.6) overnight at 4°C. Rabbit anti-TrkB [1:1,000 dilution] (Abcam #18987); mouse anti-FLAG [1:500 dilution] (Sigma-Aldrich #F3165); mouse anti-α-Tubulin [1:5,000 dilution] (Sigma-Aldrich #T9026); mouse anti-GAPDH [1:1,000 dilution] (Abcam #ab8245); rabbit anti-caveolin-2 [1:500 dilution] (Abcam #ab3417). Protein bands were detected with horseradish peroxidase (HRP)-conjugated rabbit or mouse secondary antibody [1:1,000 dilution] (Promega). The chemiluminescence signal was imaged using an Odyssey XF Imager (LI-COR) and signal intensity was quantified using ImageJ (NIH).

### Live cell imaging using molecular probes

Fluorescent probe solutions [AlexaFluor™ 488 wheat germ agglutinin (WGA) conjugate (Invitrogen #11261); AlexaFluor™ 555 cholera toxin subunit B (recombinant) (Invitrogen #C22843)] of 1mg/mL (Molecular Probes®, ThermoFisher Scientific) were prepared by dissolving 5mg of lyophilized conjugate in 5mL sterile PBS to create 1mg/mL stock solutions. Stock solutions were diluted in MEM minus phenol red (for labelling of HEK293 cells) or neurobasal medium minus phenol red (for labelling of primary neurons) to create working concentrations of 5µg/mL. Cells were seeded at densities of 1×10^4^ cells per well of Greiner 96-well plates and cultured for 2 days prior to imaging. Media was removed and 100μL labelling solution was added to cell-containing wells and incubated for 45 minutes at 37°C, 5% CO_2_. Cells were incubated with a nuclear counterstain (Hoechst 33342) (1μg/mL in appropriate medium) for 5 minutes at 37°C, 5% CO_2_. The labelling solution was removed, cells were washed 2x in PBS, and incubated in 200µL of appropriate phenol red-free media for confocal imaging. Live imaging was performed using an Opera Phenix™ High-Content Screening System (PerkinElmer) at 37°C, 5% CO_2_. Cells were visualised within a high-resolution z-stack consisting of images at 0.5µm intervals through the entire nuclear volume of the cell. Images were analysed using Harmony High-Content Imaging and Analysis Software (PerkinElmer).

### Immunocytochemistry

Cells were fixed in 4% paraformaldehyde (PFA) for 20 minutes at RT. Cells were washed 2x with PBS and blocked using 5% normal goat serum (NGS) in PBS (v/v) for 45 minutes at room temperature (RT). Cells were incubated in 0.01% Triton™ X-100 (Sigma-Aldrich) for 30 minutes at RT, then incubated in primary antibody (+ 5% NGS [v/v]) for 1 hour at RT. Rabbit anti-GM130 [1:250 dilution] (Abcam #52649); mouse anti-TGN46 [1:500 dilution] (Abcam #50595). Cells were washed 3x in PBS, then incubated in fluorescent secondary antibody (+ 5% NGS [v/v]) in a dark environment for 1 hour at RT followed by a nuclear counterstain (Hoechst 33342) (1μg/mL) for a further 5 minutes at RT. Cells were washed 3x with PBS and imaged using either an Opera Phenix™ High Content Screening System (PerkinElmer) or a Leica SP5 Confocal Microscope. Image analysis was performed using FIJI.

## Acknowledgements

We acknowledge support from a Kingsland fellowship (T.M.) and the NIHR Sheffield Biomedical Research Centre for Translational Neuroscience (IS-BRC-1215-20017). This work was supported by the Wellcome Trust (216596/Z/19/Z to J.C.-K.), and NIHR (NF-SI-0617-10077 to P.J.S.). The graphical abstract was created with BioRender.com.

## Author contributions

TM, JCK and PJS conceived and designed the study. TM, EG, AU, AH, JCK and PJS were responsible for data acquisition. TM, EG, AH and JCK were responsible for analysis of data. TM, EG, NS, RR, AH, JCK and PJS were responsible for interpretation of data. SW and BH edited the manuscript and provided input on MLR isolation. TM and JCK prepared the manuscript with assistance from all authors. All authors meet the four ICMJE authorship criteria, and were responsible for revising the manuscript, approving the final version for publication, and for the accuracy and integrity of the work.

## Declaration of interests

No authors have competing interests to declare.

## Notes

### Competing Interest Statement

The authors have declared no competing interest.

